# ERM proteins regulate the shape and number of Endoplasmic Reticulum–Plasma Membrane Junctions in neurons

**DOI:** 10.1101/2025.06.18.660273

**Authors:** Huichao Deng, Jinbo Cheng, Richard D. Fetter, Guanghui Qin, Jianxiu Zhang, Xing Liang, Caitlin A Taylor, Mingjie Zhang, Xiandeng Wu, Kang Shen

**Affiliations:** Howard Hughes Medical Institute, Department of Biology, Stanford University, Stanford, CA, USA; School of Life Sciences, Southern University of Science and Technology, Shenzhen 518055, China; Department of Biochemistry and Biophysics, UC San Francisco, San Francisco, CA, 94158, USA; Department of Health Futures, Microsoft, Redmond, WA, USA; Department of Molecular and Cellular Physiology, Stanford University School of Medicine, Stanford, CA 94305, USA; Department of Biology, Stanford University, Stanford, CA 94305, USA; Knight Initiative of Brain Resilience, WuTsai Neuroscience Institute, Stanford University, Stanford, CA, USA

## Abstract

Endoplasmic Reticulum (ER) - Plasma Membrane (PM) Junctions (EPJs) are specialized contact sites between ER membrane and the inner leaflet of PM. These junctions are critical for lipid exchange and Ca^2+^ signaling. In excitable cells like neurons and muscle, EPJs further modulate membrane excitability by regulating Ca^2+^ homeostasis. The mechanisms controlling EPJ abundance and morphology remain poorly understood. Using *in vivo* fluorescence imaging and electron microscopy of *C. elegans* neurons, we showed that EPJs form discrete, patch-like structures distributed across the soma. Through a forward genetics screen, we identified two conserved ERM (Ezrin-Radixin-Moesin) proteins, FRM-4 and FRM-1, as key regulators of EPJ shape and abundance. Both proteins localize to EPJs and exhibit liquid-liquid phase separation properties (LLPS). *in vitro*, purified FRM-4 binds to FRM-1, and together bundle filamentous actin. However, their presence in LLPS condensates and actin-bundling activity are mutually exclusive. *In vivo*, F-actin cables surround—but do not penetrate—EPJs, where FRM proteins are enriched as phase-separated condensates. Loss of FRM-4, FRM-1, or disruption of F-actin led to increased mobility of EPJs that fused into fewer but enlarged junctions. Together, our findings demonstrate that FRM-4 and FRM-1 control EPJ morphology by organizing peri-junctional F-actin networks, thereby restricting EPJ mobility and fusion.

## Main

Endoplasmic Reticulum - Plasma Membrane Junctions (EPJs) form where the ER and PM come into close proximity (7-30 nm), permitting communication between these two organelles^1–3^. These junctions serve critical functions in Ca^2+^ regulation, lipid exchange and signaling^2, 4, 5^. While EPJs are conserved across eukaryotes, their abundance, morphology, and functional roles vary significantly between cell types and physiological conditions^3–7^. Notably, excitable cells such as neurons possess highly specialized EPJs^3, 8^.

Emerging evidence suggests that dysregulation of ER-PM communication contributes to the pathogenesis of several neurological disorders, including amyotrophic lateral sclerosis (ALS), Parkinson’s disease, and Huntington’s disease^9–15^. Despite its importance, the molecular underpinnings of their development, maintenance, and functional modulation remain poorly understood. To investigate the morphology and function of EPJs, we utilized the *Caenorhabditis elegans* PVD sensory neuron as a model system. The PVD neuron features a highly polarized morphology with an exuberant branched dendritic arbor and an unbranched axon^16^. Here, we characterized the EPJs in PVD and other neurons; performed genetic screens to identify mutants with abnormal EPJ morphology and abundance and identified two conserved EPJ proteins, FRM-4 and FRM-1, which cooperatively regulate junction morphology through interactions with the actin network.

### Ultrastructural characterization of EPJs in the PVD soma reveals abundant EPJs

To understand the ultrastructure of the PVD neuron, we used serial section transmission electron microscopy (ssTEM) to generate 3D reconstructions of the PVD soma. The reconstruction revealed that the PVD soma lacks the conventional stacked ER cisternae but exhibited a simplified architecture: the ER formed a continuous, barrel-like structure in the cell cortex (Fig. 1a). The surface of ER is decorated with ribosomes, typical of rough ER (Fig. 1a, a.2). Importantly, a significant portion of the ER membrane runs close to the PM (Fig. 1a, a.1). Two distinct features indicate ER in these regions form EPJs. First, ribosomes are stringently excluded from this portion of the ER facing the PM (Fig. 1a, 1a.1, 1c). Second, the average distance between the ER and PM at these sites was around 23 nm (Fig. 1c), consistent with EPJs reported in other system^3, 6, 7^. Additionally, the 3D reconstruction of the PVD soma revealed that these EPJs are organized into discrete, patch-like domains distributed across the cell periphery, collectively resembling a soccer ball like pattern (Fig. 1b). Quantitative analysis showed that these junctions occupy approximately 18.7% of the total PM surface area. We also performed ssTEM reconstructions of additional neuronal somas in the head ganglions, which also revealed a soccer ball-like pattern of the EPJs (Extended Data Fig. 1a).

**Fig. 1.**
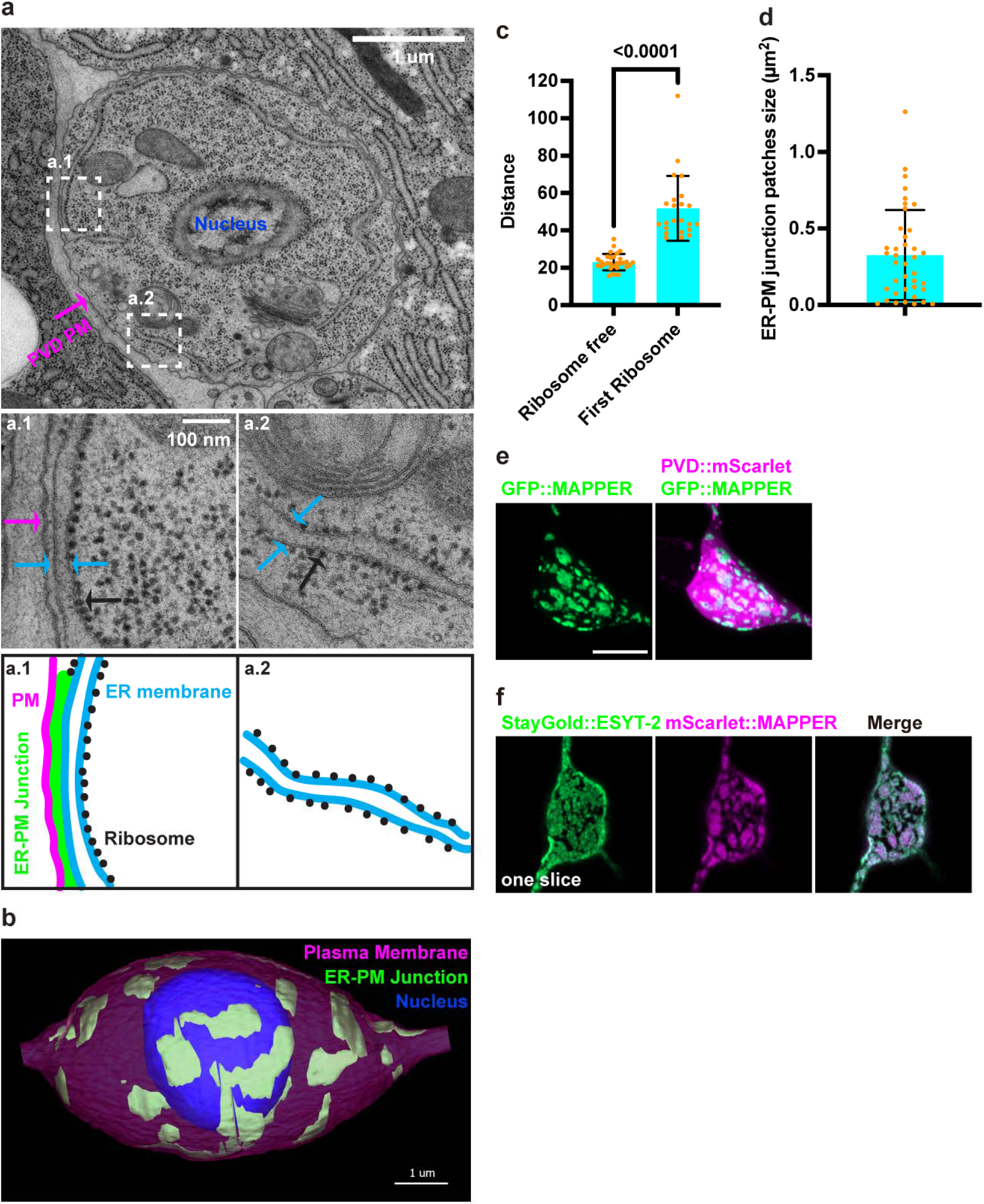
Ultrastructural characterization of EPJs in the PVD soma. a, Section of an L4 C. elegans PVD soma near the center of the cell showing the encircling rough ER with regions of EPJs. a.1) EPJ detail. a.2) Rough ER detail. b, Three-dimensional EM reconstruction of a PVD soma. c, Quantification the gap distance of EPJs and distance from first ribosome to PM outside EPJs in PVD soma according to EM. d, Quantification the EPJ patches size in PVD soma according to EM. e, Representative confocal z-stack images showing MAPPER pattern in PVD soma, visualized by ser-2Prom3::GFP::MAPPER. PVD visualized by ser-2Prom3::mScarlet. Scale bars, 5 μm. f, Representative confocal images of an PVD co-expressing ser-2Prom3::mScarlet::MAPPER and single copy des-2ps::StayGold::ESYT-2. Scale bar, 5 µm.

To enable molecular characterization of EPJs, we expressed the artificial ER-PM tether MAPPER^17^ fused to GFP (GFP::MAPPER) in PVD, which produced a GFP pattern in the soma that closely mirrored the EPJ pattern in the EM reconstruction. In PVD::GFP::MAPPER animals, the GFP signal was exclusively concentrated within the soma and displayed a patch like pattern (Fig. 1e). To validate the specificity of MAPPER as an EPJ marker in PVD, we expressed the known EPJ protein ESYT-2 tagged with StayGold (StayGold::ESYT-2) as a single copy transgene^18^. mScarlet::MAPPER and StayGold::ESYT-2 showed near complete colocalization in the PVD soma (Fig. 1f), confirming the reliability of MAPPER as a reporter of EPJs.

### FRM-4 localizes to EPJ, and regulates their morphology in a cell-autonomous manner

To uncover novel regulators of EPJs, we performed a forward genetic screen using PVD::GFP::MAPPER. From this screen, we identified the mutant allele *wy1931*, which exhibited significantly larger and fewer EPJ patches in the PVD soma (Fig. 2a, c). SNP mapping combined with whole-genome sequencing revealed that *wy1931* harbors a premature stop codon in the *frm-4* gene. To confirm that the EPJs’ phenotype was attributable to *frm-4*, we generated a null allele via CRISPR-Cas9-mediated deletion of the entire *frm-4* coding region. Both *wy1931* and *frm-4* (*null*) mutants displayed similar EPJ defects: fewer but enlarged MAPPER-labeled junctions, demonstrating that *frm-4* is essential for EPJ morphology (Fig. 2a, c). Reintroduction of wild type FRM-4 under a PVD specific promoter rescued the phenotype, indicating that *frm-4* functions cell-autonomously in neurons to regulate the number and shape of EPJs (Fig. 2a, c).

**Fig. 2.**
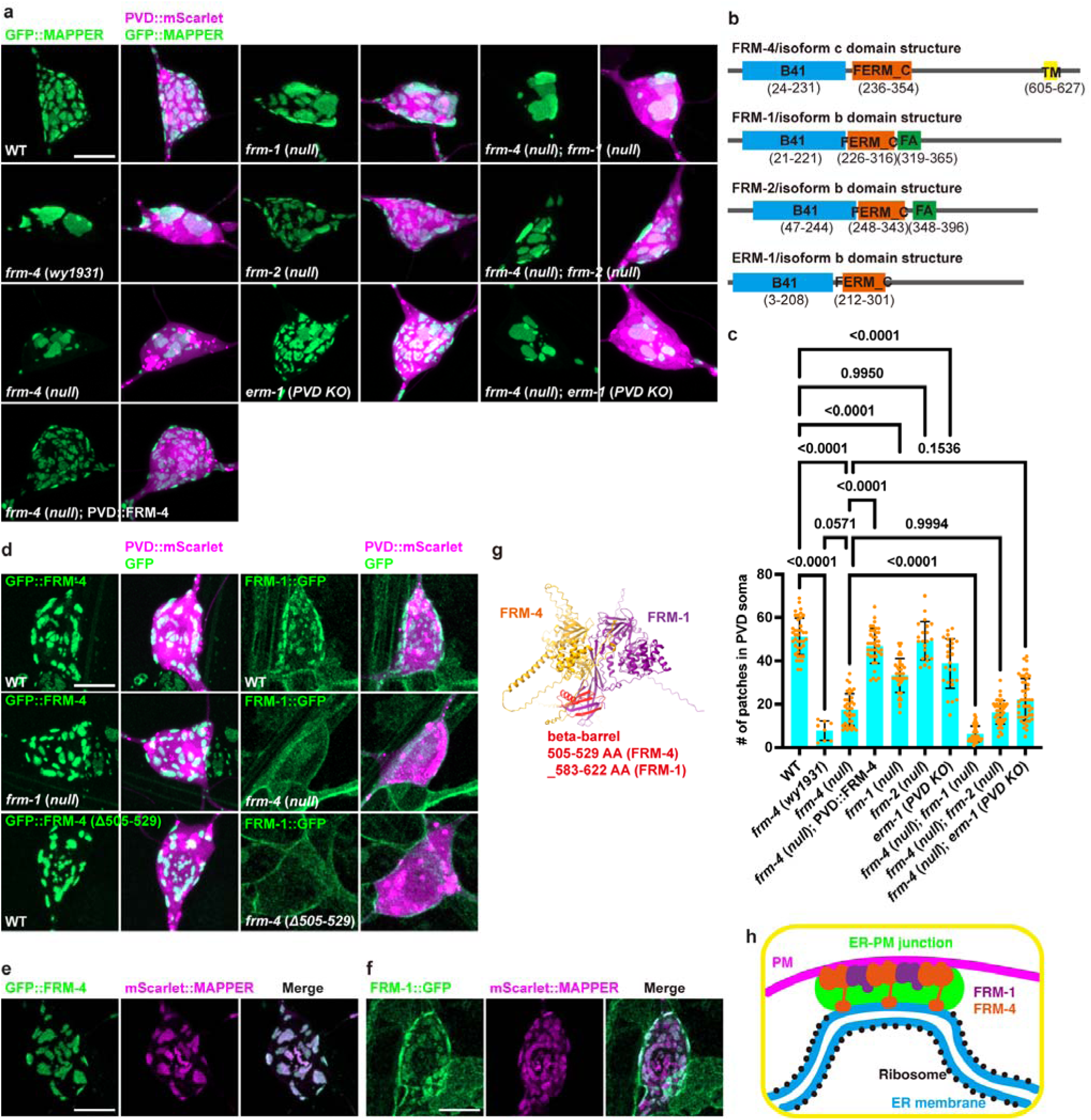
FRM-4 and FRM-1 cooperatively regulate EPJs morphology. a, Representative confocal z-stack images showing EPJs pattern in PVD soma in wild type and mutants, visualized by ser-2Prom3::GFP::MAPPER. PVD visualized by ser-2Prom3::mScarlet. Scale bar, 5 µm. b, Predicted domain structures of FRM-4, FRM-1, FRM-2, ERM-1. c, Quantification of the number of GFP::MAPPER patches in PVD soma in wild type and mutants. Data are mean ± 95% CI from independent animals. Comparisons using one-way analysis of variance (ANOVA) and Tukey’s tests. d, Representative confocal z-stack images showing endogenous GFP::FRM-4 and endogenous GFP::FRM-1 in PVD soma in wild type and mutants. PVD visualized by des-2ps::mScarlet. Scale bar, 5 µm. e, Representative confocal z-stack images of an PVD co-expressing ser-2Prom3::mScarlet::MAPPER and endogenous GFP::FRM-4. Scale bar, 5 µm. f, Representative confocal z-stack images of an PVD co-expressing ser-2Prom3::mScarlet::MAPPER and endogenous GFP::FRM-1. Scale bar, 5 µm. g, Structure prediction of interaction between FRM-4 and FRM-1. h, A schematic model for FRM-4 and FRM-1 at a EPJ.

To determine the subcellular localization of FRM-4, we inserted a conditional GFP cassette in frame into the endogenous *frm-4* locus and used a PVD specific FLP recombinase to achieve cell-specific endogenous tagging^19^. GFP::FRM-4 displayed a patch-like pattern (Fig. 2d) and showed near complete colocalization with mScarlet::MAPPER (Fig. 2e), indicating that FRM-4 localizes predominantly to EPJs.

FRM-4 contains a B41 (Band 4.1) and a FERM_C (Four-point-one, Ezrin, Radixin, Moesin) domain followed by a transmembrane (TM) domain with no N-terminal signal peptide (Extended Data Fig. 2a). B41 and FERM_C domains are found in ERM family of proteins which is known to link actin networks to the plasma membrane^20^. Based on the domain structures, we propose that FRM-4 is a transmembrane ER protein which is strongly enriched at the EPJs. To dissect the function of each domain, we generated three endogenous deletion alleles: *frm-4* (*ΔB41*), *frm-4* (*ΔFERM_C*), and *frm-4* (*ΔTM*) using CRISPR-Cas9 (Extended Data Fig. 2a, b). Each of these mutants phenocopied the *frm-4* (*null*) allele, exhibiting enlarged and sparse EPJs (Extended Data Fig. 2b, c). Thus all three domains are required for FRM-4’s function.

To further understand the role of each domain in FRM-4 localization, we examined the subcellular distribution of the corresponding endogenous GFP-tagged deletion proteins. All three domain deletions led to greatly decreased GFP signal (Extended Data Fig. 2d, e), suggesting that the B41, FERM_C, and TM domains are all required for FRM-4 protein stabilization. Upon signal overexposure, endogenous GFP::FRM-4 (ΔB41) displayed an ER-like pattern (Extended Data Fig. 2d, e, f), indicating that the B41 domain might contribute to bring the ER membrane to the PM. In contrast, endogenous GFP::FRM-4 (ΔFERM_C) exhibited negligible signal, underscoring the FERM_C domain’s critical role in protein stability (Extended Data Fig. 2d, e, f). Interestingly, endogenous GFP::FRM-4 (ΔTM) showed a plasma membrane associated pattern, suggesting that the TM domain anchors FRM-4 to the ER membrane (Extended Data Fig. 2d, e, f).

To assess organism-wide expression, we inserted a StayGold fluorescence protein in frame into the endogenous *frm-4* locus. This knock in strain revealed broad FRM-4 expression across various tissues including many neurons, intestine, hypodermis, muscle, germline, oocytes, spermatheca and embryos (Extended Data Fig. 2g, h), implying potential roles for FRM-4 in multiple cell types. In non-neuronal tissues, FRM-4 displayed as puncta of various density and regularity. In neurons, FRM-4 showed patch-like distribution similar to the pattern in PVD, suggesting that neurons have specialized FRM-4 subcellular pattern. Example images are show for FRM-4::staygold in the FLP and HSN neurons (Extended Data Fig. 1g).

### FRM-4 and FRM-1 cooperatively regulate EPJs’ morphology

Based on sequence homology, *frm-4* has three paralogs in *C. elegans*: *frm-1*, *frm-2*, and *erm-1*, all of which share conserved B41 and FERM_C domains. However, unlike *frm-4*, none of these paralogs possess predicted transmembrane domains (Fig. 2b). To assess their functional relevance in EPJ regulation, we generated complete gene deletion alleles of *frm-1* and *frm-2*, using CRISPR-Cas9. *erm-1* is an essential gene, therefore we created a PVD cell specific knockout, *erm-1* (*PVD KO*) to assess its function. Among them, only the *frm-1* (*null*) mutant exhibited a notable reduction in the number of EPJ patches in PVD, though this phenotype was milder compared to the *frm-4* (*null*) mutant (Fig. 2a, c). Notably, the *frm-4* (*null*); *frm-1* (*null*) double mutant displayed a significantly enhanced phenotype relative to the *frm-4* (*null*) single mutant (Fig. 2a, c), indicating that *frm-1* and *frm-4* function cooperatively in EPJ organization, with *frm-4* playing a more important role. The *frm-4* (*null*) *frm-2* (*null*) or *frm-4* (*null*) *erm-1* (*PVD KO*) did not enhance the *frm-4* (*null*) single mutant phenotype (Fig. 2a-c).

To further characterize FRM-1’s role, we examined its subcellular localization. Endogenously tagged FRM-1::GFP exhibited a patch-like pattern (Fig. 2d), which largely colocalized with the EPJ marker mScarlet::MAPPER (Fig. 2f). This colocalization suggests that FRM-1 predominantly localizes to EPJs, consistent with the EPJ defects observed in the *frm-1* (*null*) mutant. Lacking a transmembrane domain, FRM-1 is predicted to be a cytosolic protein. We next asked how it localizes to EPJs. Given FRM-4’s localization and function at EPJs, we tested the localization of endogenous FRM-1::GFP in the *frm-4* (*null*) background and observed FRM-1 completely loses its enrichment at EPJs and becomes diffusely localized near plasma membrane (Fig. 2d). This result indicates that FRM-4 is required for recruiting FRM-1 to EPJs. In the *frm-4* mutants, while EPJs become fewer and enlarged (Fig.2a-c), FRM-1 is no longer enriched at the EPJs. To ask whether FRM-4 and FRM-1 can directly interact with each other, we used AlphaFold 3 to predict the structures of these two proteins. The prediction indicated a high likelihood of interaction between FRM-1 and FRM-4 with the FRM-4 amino acids 505 to 529 forming a β-barrel and interacting with amino acid 583-622 of FRM-1 (Fig. 2g). We generated an endogenous deletion mutant of FRM-4 lacking amino acids from 505 to 529 (Fig. 2d, g) and tested how this allele affects EJP morphology and FRM-1 localization. While endogenous GFP::FRM-4 (Δ505-529) retained the typical EPJ localization pattern (Fig. 2d), endogenous FRM-1::GFP in *frm-4* (*Δ505-529*) background lost junctional enrichment and becomes diffusely localized to plasma membrane (Fig. 2d). These data strongly suggest that FRM-1 is recruited to EPJs by interacting with FRM-4.

To directly probe the interaction between FRM-1 and FRM-4, we biochemically purified FRM-1 full-length protein and FRM-4 near full-length protein (with C-terminal transmembrane truncated, denoted as FRM-4 herein). We performed size exclusion chromatography coupled with multiangle light scattering (SEC-MALS) analysis to measure the assembly states of FRM-1 and FRM-4. Increasing concentration of FRM-1 or FRM-4 proteins leads to subtle shift towards smaller elution volume, indicating that both FRM-1 and FRM-4 might form weak homo-oligomers (Extended Data Fig. 3a, b). When mixed together, there was a complex peak located at a smaller elution volume compared to either one (Extended Data Fig. 3c), suggesting FRM-1 and FRM-4 bind to each other, likely with low affinity. In parallel, GST pull-down experiment also suggest that FRM-4 and FRM-1 exhibit substoichiometric interactions with each other (Extended Data Fig. 3d).

To determine the contribution of individual domains to FRM-1 protein stability, we analyzed the subcellular localization and fluorescence intensity of endogenously GFP-tagged FRM-1 deletion variants. Deletion of either the B41 domain or the FERM_C domain resulted in markedly reduced GFP signal (Extended Data Fig. 4a, b, c), indicating that both domains are critical for stabilizing the FRM-1 protein. This requirement mirrors that of FRM-4, which similarly depends on these conserved domains. In contrast, deletion of the FA domain did not affect GFP intensity or localization, suggesting that the FA domain is dispensable for FRM-1 stability (Extended Data Fig. 4a, b, c).

### EPJs exhibit increased dynamics and fusion in *frm-4*; *frm-1* double mutants

To elucidate the mechanisms underlying the observed enlargement and reduced number of EPJs in *frm-4*; *frm-1* double mutants, we analyzed the temporal dynamics of EPJ morphology during development. Specifically, we quantified the number of EPJs from the L4 larval stage to day 1 of adulthood (24 hours post-L4). In wild type animals, the number of EPJ remained relatively stable over this period (Fig. 3a, b). In contrast, *frm-4*; *frm-1* double mutants exhibited reduction in patch number during development (Fig. 3a, b). Notably, at earlier developmental stages, the double mutants showed both large and smaller EPJ patches with reduced number of patches compared to the wild type L4 animals (Fig. 3a, c). When the mutant animals reached adult stage, most only large patches remain with very low number of EPJs (Fig. 3a-d). These findings imply that EPJs might fuse with each other in the absence of FRM-4 and FRM-1.

**Fig. 3.**
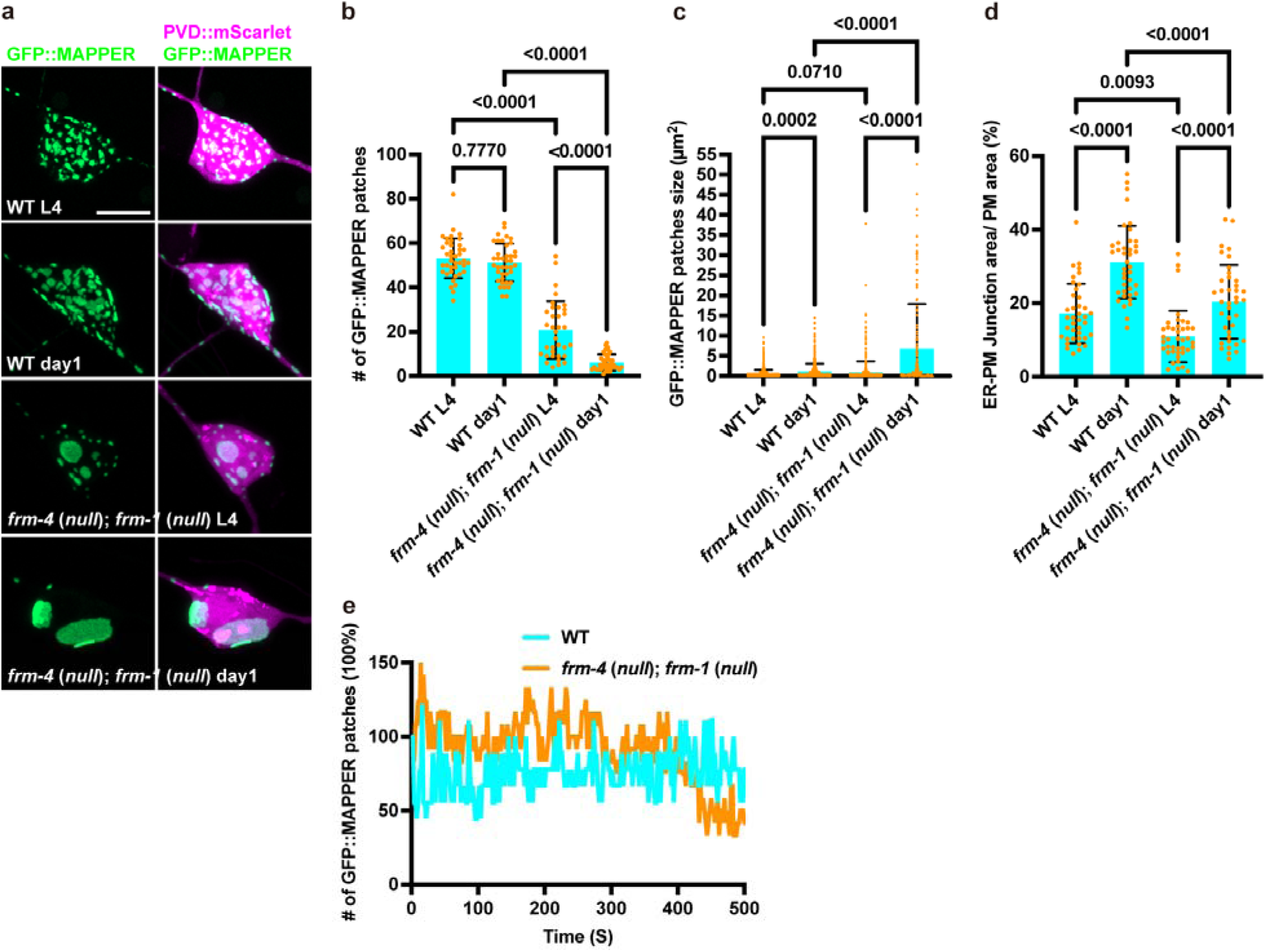
EPJs exhibit increased dynamics and fusion in *frm-4*; *frm-1* double mutants during development. a, Representative confocal z-stack images showing EPJ pattern in PVD soma in wild type and mutants, visualized by ser-2Prom3::GFP::MAPPER. PVD visualized by ser-2Prom3::mScarlet. Scale bar, 5 µm. b, Quantification of the number GFP::MAPPER patches in PVD soma in wild type and mutants. Data are mean ± 95% CI from independent animals. Comparisons using one-way analysis of variance (ANOVA) and Tukey’s tests. c, Quantification of the size of GFP::MAPPER patches in PVD soma in wild type and mutants. Data are mean ± 95% CI from independent animals. Comparisons using one-way analysis of variance (ANOVA) and Tukey’s tests. d, Quantification the EPJ area/PM area in PVD soma in wild type and mutants. Data are mean ± 95% CI from independent animals. Comparisons using one-way analysis of variance (ANOVA) and Tukey’s tests. e, Time-lapse analyses of the number of GFP::MAPPER patches in PVD soma in wild type and mutants.

To directly assess EPJ dynamics, we performed time-lapse imaging of EPJs at the L2 larval stage. Compared with wild type animals, EPJ patches in *frm-4*; *frm-1* double mutants displayed significantly greater motility and increased frequency of fusion events over time (Movie 1 and Fig. 3e). These results support the hypothesis that FRM-4 and FRM-1 act to stabilize EPJs during development, limiting their mobility and preventing excessive fusion.

### F-actin is required for EPJ and FRM-4 morphology

How do FRM-4 and FRM-1 restrict EJP mobility and prevent excessive fusion? FRM-4 and FRM-1 are members of the evolutionarily conserved ERM protein family, characterized by a common phosphatidylinositol 4,5-bisphosphate (PIP2)-binding FERM domain and a C-terminal F-actin-binding site. These domains enable ERM proteins to act as molecular linkers between the plasma membrane and the actin cytoskeleton^20^. We hypothesized that FRM-4 and FRM-1 stabilize EPJs through binding with actin cytoskeleton.

To directly assess the role of F-actin in EPJ morphology, we employed the actin-depolymerizing toxin SpvB, an ADP-ribosyltransferase that inactivates actin monomers by ADP-ribosylation of a conserved arginine residue on actin monomer, thereby preventing polymerization^21^. We expressed SpvB specifically in PVD neurons using the PVD::SpvB transgene^21^ in GFP::MAPPER strain. Disruption of F-actin in PVD neurons led to a marked reduction in the number of EPJs within the PVD soma (Fig. 4a, b). This phenotype resembled that observed in *frm-4*; *frm-1* double mutants, suggesting a functional link between actin integrity and FRM-mediated junction stability. Consistently, expression of PVD::SpvB in an endogenous GFP::FRM-4 background significantly reduced the number of visible FRM-4 patches at the plasma membrane (Fig. 4c, d). Together, these results demonstrate that F-actin is critical for maintaining both EPJ morphology and suggest that FRM-4 and FRM-1 function as cytoskeletal tethers to restrict EPJ mobility and prevent EPJ fusion.

**Fig. 4.**
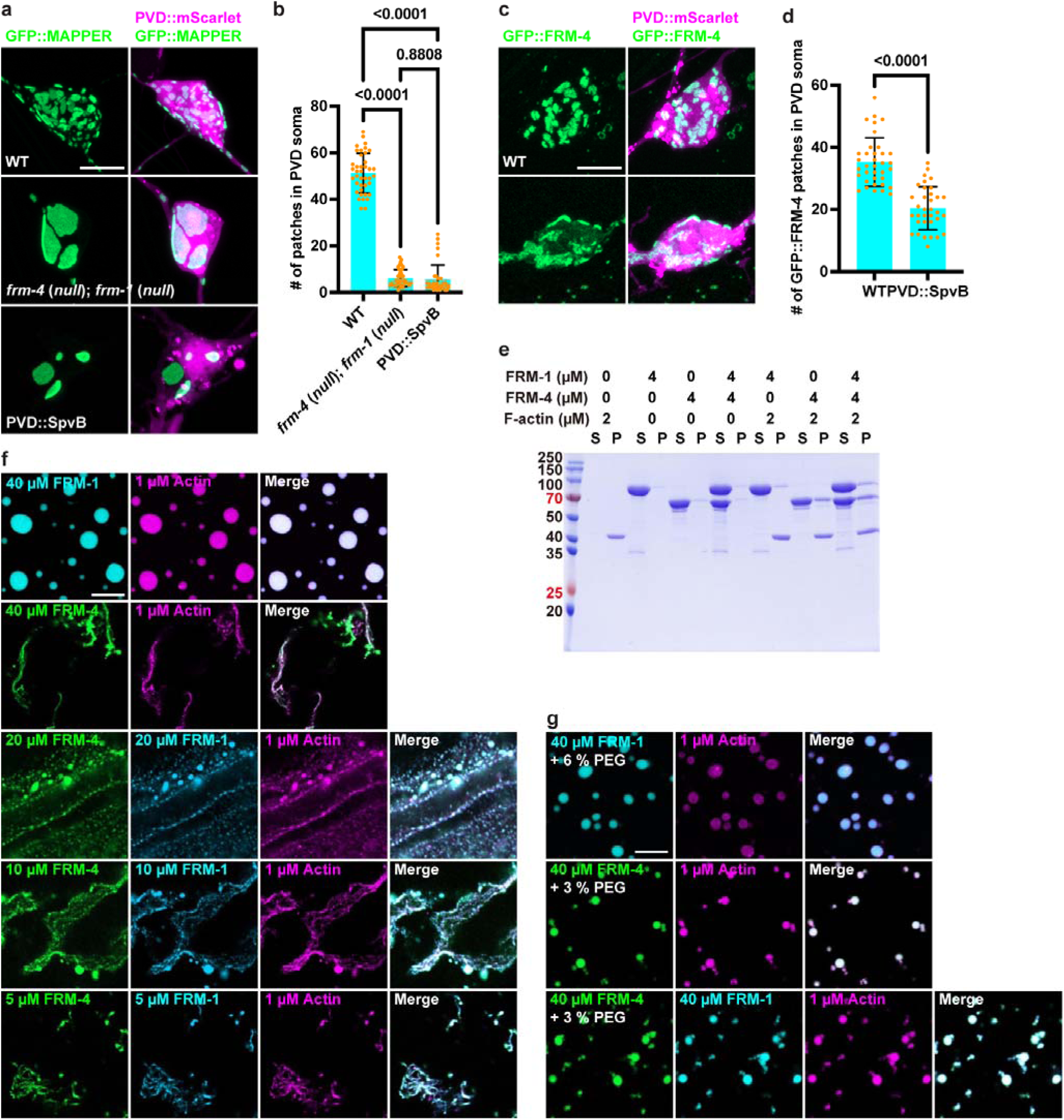
F-actin is required for EPJ and FRM-4 morphology. a, Representative confocal z-stack images showing EPJ pattern in PVD soma in wild type and mutants, visualized by ser-2Prom3::GFP::MAPPER. PVD visualized by *ser-2Prom3::mScarlet*. Scale bar, 5 µm. b, Quantification of the number of GFP::MAPPER patches in PVD soma in wild type and mutants. Data are mean ± 95% CI from independent animals. Comparisons using one-way analysis of variance (ANOVA) and Tukey’s tests. c, Representative confocal z-stack images showing endogenous GFP::FRM-4 in PVD soma in wild type and mutant. PVD visualized by des-2ps::mScarlet. Scale bar, 5 µm. d, Quantification of the number of endogenous GFP::FRM-4 patches in PVD soma in wild type and mutant. Comparisons by two-tailed t-test. e, Co-sedimentation assay to test the interaction between FRM-4, FRM-1 and F-actin. f, When FRM proteins were incubated with G-actin in the polymerization buffer prior to phase separation induction (by adding 3% PEG), FRM-4 but not FRM-1 could efficiently bundle actin. The addition of FRM-1 could lower the threshold of FRM-4 phase separation and enhance actin bundling efficiency. Scale bar, 10 µm. g, Preformed FRM-1 and/or FRM-4 condensates barely bundle actin. Scale bar, 10 µm.

### FRM-4 bundle actin filaments *in vitro*

To examine whether FRM-4 and FRM-1 directly bind to F-actin, we performed a co-sedimentation assay. At high-speed centrifugation, F-actin is present in the pellet and depleted from the supernatant. However, purified FRM-1 or FRM-4 remains in the supernatant (Fig. 4e). Upon incubation with F-actin, FRM-4 becomes present in the pellet fraction, suggesting direct binding between F-actin and FRM-4. In contrast, FRM-1 does not interact with F-actin in this assay. Incubation of both FRM-4 and FRM-1 led to both protein co-pellet with F-actin (Fig.4e).

Next, we asked how FRM-4/FRM-1 interact with F-actin. To address this, we first tackled FRM-4 and FRM-1 assembly mechanism. We found that both purified FRM-4 and FRM-1 undergo phase separation in the presence of the crowding reagent, PEG8000 (Extended Data Fig. 5a, b). Upon mixing, FRM-4 and FRM-1 mutually promote each other to phase separate (Extended Data Fig. 5c). This is reminiscent of the weak oligomerization of FRM-4 and FRM-1 detected by gel filtration (Extended Data Fig. 3a-c). It is conceivable that the high protein concentration in the condensates induces the oligomerization states, which in return promotes phase separation.

Next, we investigated F-actin bundling property of FRM-4/FRM-1. We pre-incubated G-actin and FRM-4 or FRM-1 proteins at the desired concentration in actin polymerization buffer to allow F-actin formation, then added PEG8000 to trigger the phase separation, immediately followed by imaging experiments. We found that FRM-1 and G-actin are both present in the same phase condensate droplets. Interestingly, FRM-4 colocalizes with bundled F-actin filaments (Fig. 4f). The thick F-actin staining FRM-4 confirms that FRM-4 can directly bind and bundle F-actin, while FRM-1 alone does not bundle to F-actin. Furthermore, adding FRM-1 to FRM-4 together to F-actin lead to the F-actin bundling and colocalization of both proteins with F-actin, even at lower concentration of the FRM proteins (Fig. 4f). These *in vitro* result mirrors the mutant phenotypes, which show a central role of FRM-4 and synergistic function between FRM-1 and FRM-4 in controlling the EPJ morphology.

To further understand the relationship between the phase separation and the F-actin bundling activities of FRM-1 and FRM-4, we mixed all the components (FRM-4/FRM1, G-actin, and polymerization buffer) at the same time. We found that neither FRM-4, FRM-1 alone, nor together can bundle F-actin (Fig. 4g). Instead, the form phase droplets which also recruits G-actin. These results suggest that FRM-4’s ability to bundle to F-actin and form phase condensates are mutually exclusive. Together, these results show that FRM-4 and FRM-1 lose their actin bundling activities when they form liquid-liquid phase condensates.

### FRM-4 promotes local assembly of actin filaments *in vivo*

To further test whether FRM-4 organizes actin filament *in vivo*, we first examined the spatial relationship between FRM-4 and actin filaments using the filamentous actin marker GFP::moesinABD^22^ in animals expressing endogenous mScarlet::FRM-4. GFP::moesinABD labeled F-actin filaments form cable-like structures surrounding the mScarlet::FRM-4 patches (Fig. 5a), suggesting that actin filaments surround the EPJ patches.

**Fig. 5.**
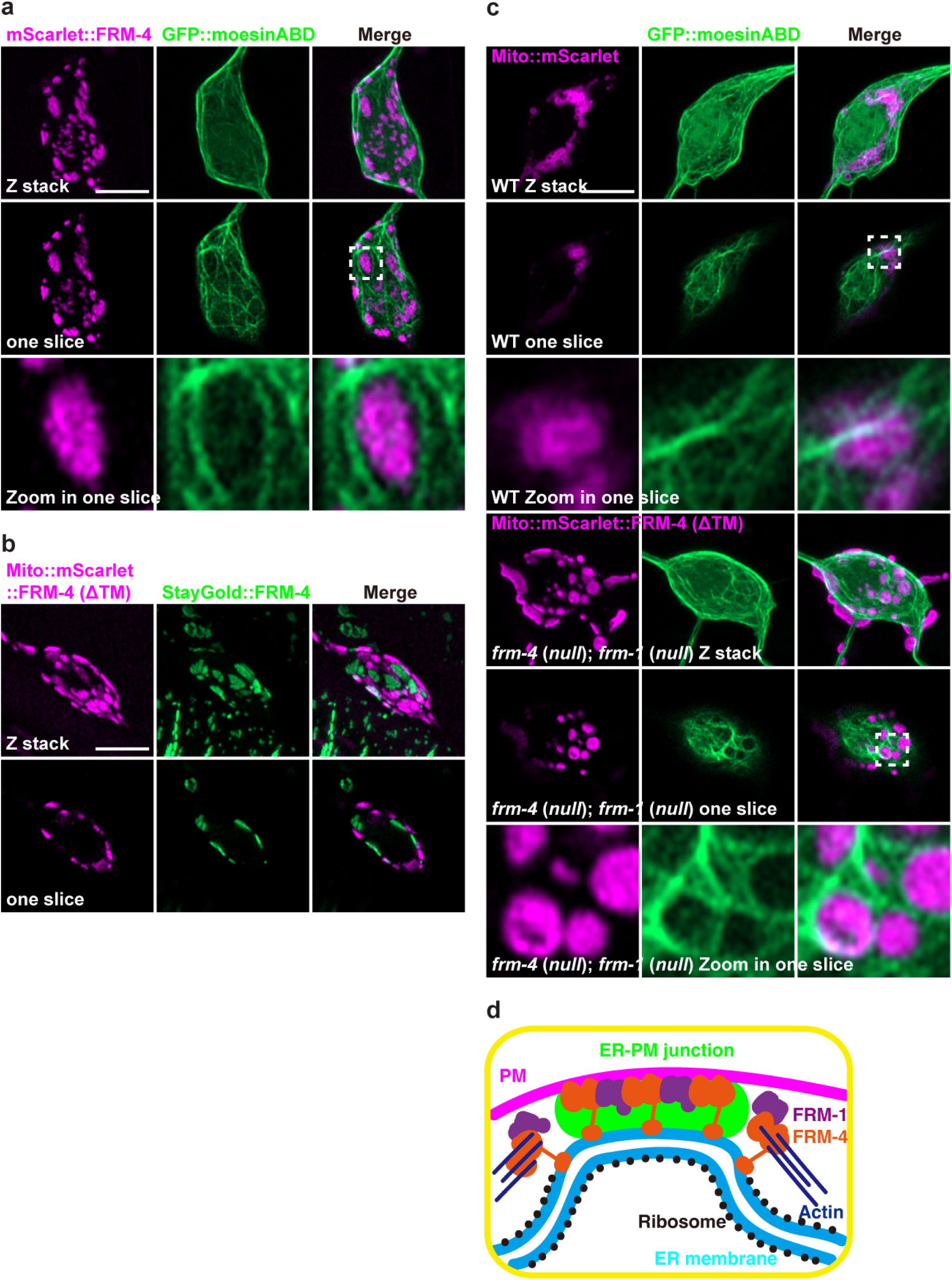
FRM-4 promotes local assembly of actin filaments *in vivo*. a, Representative confocal z-stack images of an PVD co-expressing ser-2Prom3::GFP::moesinABD and endogenous mScarlet::FRM-4. Scale bar, 5 µm. b, Representative confocal images of an PVD co-expressing ser-2Prom3::Mit(TOMM-20)::mScarlet::FRM-4 (ΔTM) and endogenous StayGold::FRM-4. Scale bar, 5 µm. c, Representative confocal images of an PVD co-expressing ser-2Prom3::Mito(TOMM-20)::mScarlet and ser-2Prom3::GFP::moesinABD in wild type. ser-2Prom3::Mito(TOMM-20)::mScarlet::FRM-4(ΔTM) and ser-2Prom3::GFP::moesinABD in *frm-4*; *frm-1* double mutans. Scale bar, 5 µm. d, A schematic model of FRM-4 and FRM-1’s phase condensates at EPJ and their actin bundling activity at peri-junctional ER.

To further investigate whether FRM-4 is sufficient to recruit or organize actin filaments, we redirected a truncated version of FRM-4 lacking the transmembrane domain, FRM-4(ΔTM) to the outer mitochondrial membrane by fusing it with the mitochondrial targeting sequence from TOMM-20. Interestingly, expression of Mito (TOMM-20)::mScarlet::FRM-4 (ΔTM) in neurons led to the striking recruitment of mitochondria to the close to the plasma membrane (Fig. 5b). The mitochondria next to PM forms patch-like structures that are in between the endogenous FRM-4 labeled EPJs (fig. 5b). This result indicates that FRM-4 is sufficient to recruit mitochondria membrane to the plasma membrane. Next, we asked whether FRM-4 can bundle F-actin around mitochondria. Wild type mitochondria are not consistently surrounded by F-actin. However, expression of Mito (TOMM-20)::mScarlet::FRM-4 (ΔTM) in the *frm-4*; *frm-1* double mutant background caused striking changes of F-actin distribution, which tightly surround mitochondria (Fig. 5c), further indicating that FRM-4 is sufficient to bundle F-actin.

Together, these results are consistent with the following model. FRM-4 is an ER transmembrane protein with actin and plasma membrane binding activities. FRM-4 and FRM-1 form condensate at the junction using a phase separation mechanism. Within the condensate FRM-4 and FRM-1 do not show actin binding and bundling activity. The fluidic nature of the phase condensate at the EPJs allows FRM-4 proteins to diffuse out of the junctions and presumably to peri-junctional ER membrane, where it bundles actin to form an “actin cage” and restrict the lateral mobility of EPJs (Fig. 5d).

## Discussion

Previous studies showed that excitable cells such as muscles and neurons have abundant ER-PM junctions compared to non-excitable cells^23^. EPJs in mammalian neurons represent over 10% of the PM area, with larger EPJs in the soma compared to neurites^3^. In PVD, we found that the EPJs are also prominent in soma and accounts for 18% of the PM area, demonstrating the striking similarity between C. elegans and vertebrate neurons. Using MAPPER, Hsieh and colleagues showed that cortical actin is in close proximity to EPJs and that disruption of cortical actin results in increase lateral motility and decreased number of EPJs in non-neuronal cells^24^. Our *in vivo* results closely mirror these findings, suggesting that F-actin is a conserved cellular mechanism to regulate EPJs across different cell types. Further, we discovered FRM-4 and FRM-1, novel junctional proteins which link cortical actin to the junctions to stabilize the EPJs. FRM-4 and FRM-1 are both conserved proteins and their mammalian homologs might play similar functions. Mechanistically, FRM-4 and FRM-1 use phase separation mechanisms to localize at EPJs. However, there is little F-actin within the EPJs. Instead, cortical actin is present adjacent to the junctions. Interestingly, our *in vitro* reconstitution experiments showed that FRM-4 can directly bind and bundle F-actin but loses this ability when it is in a phase condensate. We propose that the fluidic exchange of FRM-4 between the phase condensates within EPJ and the surrounding ER membrane allows a “ring” of FRM-4 to bundle actin. Through this interaction, cortical actin restricts the movement of EPJs and stabilize the junctions.

## Supporting information

Supplemental Movie 1 Left, WT. Right, frm-4;frm-1 mutant.

## Acknowledgments

We thank members of the Shen lab for their scientific feedback and discussion. Funding: K. Shen is an Investigator in the Howard Hughes Medical Institute. This project is partially supported by the Knight Initiative of Brain Resilience at Stanford University.

## Author contributions

Conceptualization, H.D. and K.S.

Methodology, H.D., J.C., R.F., G.Q., J.Z., X.L., C.T., M.Z., X.W. and K.S.

Investigation, H.D., J.C., R.F., G.Q., J.Z., X.L., C.T., M.Z., X.W. and K.S.

Writing-Original Draft, H.D. X.W. and K.S.

Writing-Review & Editing, H.D., J.C., R.F., G.Q., J.Z., X.L., C.T., M.Z., X.W. and K.S.

Funding Acquisition, K.S.

Resources, H.D. and K.S.

Supervision, K.S.

## Methods

### C. elegans strains

*C. elegans* strains were grown on OP50 E. coli-seeded nematode growth media (NGM) plates at 20°C, following standard protocols^25^. N2 Bristol is the wild-type reference strain, and all the mutants were isolated from N2.

All the endogenous knock in worms were generated by CRISPR/Cas9 editing^26^. Transgenic strains were generated using standard microinjection techniques.

All the *C. elegans* strains used in this study are listed in table S1.

### Molecular biology

Plasmids, crRNAs, primers sequence used to generate transgenic or knock in *C. elegans* strains in this study are listed in table S1. Plasmids were generated using In-fusion cloning kit. crRNAs used for CRISPR/Cas9 knock in were ordered from IDT.

### Electron microscopy

Worms were prepared for conventional EM by high pressure freezing/freeze-substitution. Worms in M9 containing 20% BSA and E. coli were frozen in 100 μm well specimen carriers (Type A) opposite a hexadecane coated flat carrier (Type B) using a Leica ICE high-pressure freezer (Leica Microsystems, Deerfield, IL). Freeze-substitution in 1% OsO4, 0.1% uranyl acetate, 1% methanol in acetone, containing 3% water^27, 28^ was carried out with a Leica AFS2 unit. Following substitution, samples were rinsed in acetone, infiltrated and then polymerized in Eponate 12 resin (Ted Pella, Inc, Redding, CA). Serial 40 nm sections were cut with a Leica UC7 ultramicrotome using a Diatome diamond knife, picked up on Pioloform coated slot grids and stained with uranyl acetate and Sato’s lead^29^. Sections were imaged with an FEI Tecnai T12 TEM at 120 kV using a Gatan 4k x 4k Rio camera. TrakEM2 in Fiji was used to align serial sections^30–32^. Modeling of serial sections was performed with IMOD^33^.

### Microscopy

*C. elegans* fluorescence imaging was performed on a Zeiss LSM 980 Airyscan 2 system with a 63× Plan-Apochromat 1.4NA objective and 488 nm, or 561 nm lasers. The Airyscan 4Y multiplex mode was used to capture super resolution images. Animals were imaged live on 5% agarose pads in 10 mM levamisole for stable imaging. Animals were imaged live on 5% agarose Petri dishes (MatTek P35G-1.5-14-C) in 2.5 mM levamisole for time lapse imaging.

### Image analysis and quantification

Airyscan images and analysis were processed with Zeiss ZEN software. Statistical calculations and graphing were done in Prism 10 (GraphPad).

EPJs patch area workflow was implemented in Python with NumPy, SciPy, scikit-image, and Trimesh. First, voxels belonging to the surface patches were separated from background by global intensity thresholding with the Li method. When the patch mask was mis-registered relative to the cell surface, the cell mask was iteratively eroded until maximal overlap was achieved. A marching-cubes isosurface was then extracted from this optimally eroded cell mask, producing a triangulated shell in physical (µm) units. The cell’s surface area was obtained by summing the areas of these triangles. Intersecting the shell mesh with the patch mask and summing the areas of the intersecting faces yielded the surface area for each individual patch.

Time lapse EPJs patch segmentation and tracking were performed in Python with NumPy, SciPy, scikit-image, and Trackpy. For each video frame, Otsu’s method determined a global intensity threshold, producing a binary mask of candidate patches. Touching or overlapping patches were separated with a marker-based watershed transform. The resulting label images were then used for patch counting and for linking objects across successive frames with Trackpy to obtain complete trajectories.

### Structure prediction

The structure prediction of the FRM-4 and FRM-1 complex was performed using AlphaFold 3^34^. The resulting structural model was visualized using ChimeraX^35^.

### Protein expression and purification

FRM-1 full-length and FRM-4ΔTM genes were cloned into a modified pET32a vector, with a Trx-6xHis tag and a protease 3C cutting site at the N-terminal of the multi-clone sites. Sequences were confirmed by sequencing. Proteins were expressed in Escherichia coli BL21-CodonPlus(DE3)-RIL cells (Agilent). Cells were grown in LB at 37°C with 220 rpm shaking to an OD600 of ∼ 0.8, followed by 0.5 mM IPTG induction for protein expression. Continue to culture the cells overnight for protein expression. Collect the cell pellet by centrifuging at 4,200 rpm for 10 min at 4°C. Resuspend the pellet with binding buffer composed of 50 mM Tris pH 8.0, 1000 mM NaCl, 5 mM Imidazole, supplemented with 1 mM PMSF. Homogenize the suspension with a French press high-pressure homogenizer, followed by centrifugation at 18,000 rpm for 30 min at 4°C. The supernatant was collected and drain through a gravity Ni2+-NTA Sepharose (GE Healthcare) column, washed with binding buffer three times, and eluted with 50 mM Tris pH 8.0, 500 mM NaCl, 300 mM imidazole. Then the proteins were subject to size-exclusion chromatography with HiLoad 26/600 Superdex 200pg column, pre-equilibrated with 50 mM Tris pH 8.0, 100 mM NaCl, 1 mM EDTA, 1 mM DTT. Fractions with proper proteins were pooled and digested by 3C protease overnight at 4°C. An anion or cation exchange column was used to remove the Trx-6xHis tag and other impurities for FRM-1 and FRM-4, respectively. Then the proteins were loaded to another round of HiLoad 26/600 Superdex 200pg column equilibrated with HBS (HEPES buffer saline: 20 mM HEPES pH 7.5, 100 mM NaCl, 1 mM TCEP). Proteins were concentrated with an Amicon ultra centrifugal filter (MWCO = 10 kDa, Millipore) to ∼ 10 mg/ml and aliquoted into small fractions. Proteins were then flash frozen in liquid nitrogen and stored at -80°C.

### Size exclusion chromatography coupled with multiangle light scattering (SEC-MALS)

The SEC-MALS system is composed of a multi-angle light scattering (MALS) detector (miniDawn, Wyatt), a differential refractive index (dRI) detector (Optilab, Wyatt), and a Liquid chromatography (LC) system (AKTA pure, GE Healthcare). For each experiment, 200 μl sample was loaded via injection loop (100 μl) into a Superdex 200 10/300 GL column (GE Healthcare) equilibrated with HBS. Data were analyzed by ASTRA6 (Wyatt).

### F-actin binding

G-actin proteins (Cytoskeleton, AKL99) were reconstituted in 5 mM Tris pH 8.0, 0.2 mM CaCl2, 0.2 mM ATP, 5% (w/v) sucrose, and 1% (w/v) dextran, and stored at -80°C. For actin polymerization, a 10x polymerization buffer (10 mM ATP, 20 mM MgCl2, 20 mM DTT in HBS) was prepared. The basal buffer for F-actin formation and bundling was HBS. The 1x polymerization buffer contains 20 mM HEPES pH 7.5, 100 mM NaCl, 1 mM TCEP, 1 mM ATP, 2 mM MgCl2, 2 mM DTT. Dilute G-actin with HBS and 1x polymerization buffer to a final concentration of 20 μM. Incubate at room temperature for 1 h. Then FRM-4 or FRM-1 proteins were mixed with F-actin at the desired concentration, and incubate at room temperature for another 30 min. Centrifuge the mixture at 100,000g for 20 min at 4°C. The pellet was resuspended with HBS and analyzed together with the supernatant using SDS-PAGE.

### Actin bundling

10% Rhodamine-labelled G-actin was reconstituted as mentioned above. For actin bundling (Fig. 4f), mix FRM-4/FRM1 proteins with G-actin at the desired concentration in 1x polymerization buffer. Incubate at room temperature for 1 h. Then add 3% PEG8000 to trigger phase separation to occur. Transfer the mixture to the imaging chamber immediately, followed by fluorescence imaging. Alternatively, for Fig. 4g, FRM-4/FRM-1, G-actin, PEG8000 were simultaneously mixed in 1x polymerization buffer, incubate at room temperature for 1 h before imaging.

## Supplemental information titles and legends

Table S1 *C. elegans* strains, Plasmids, crRNAs, Primers used in this study.

---

**Extended Data Fig. 1 for Fig. 1.**
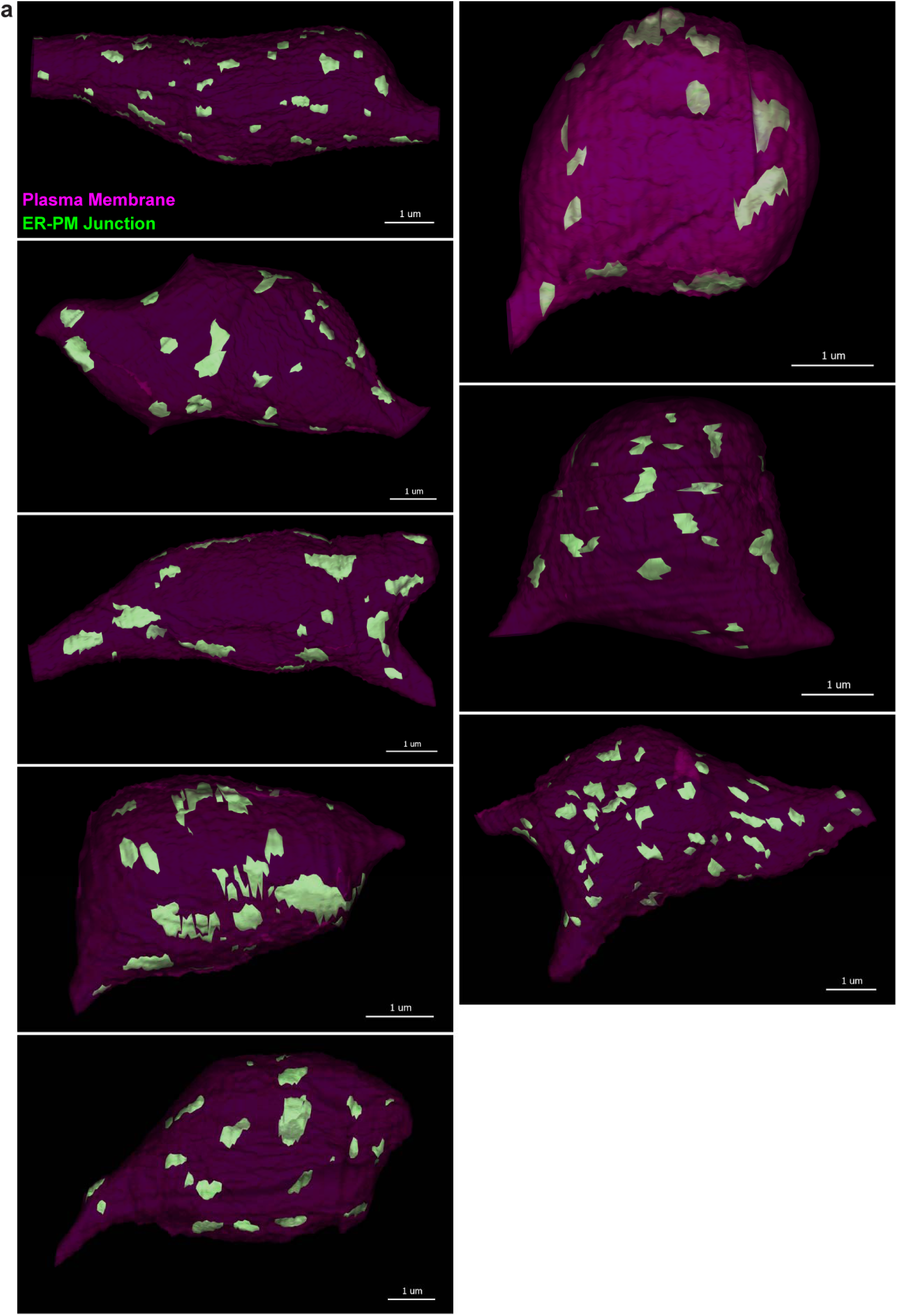
Three-dimensional EM reconstructions of head neuron somas in a Day 1 worm. The determination of ER-PM junctions is the same as in Fig. 1.

**Extended Data Fig. 2 for Fig. 2.**
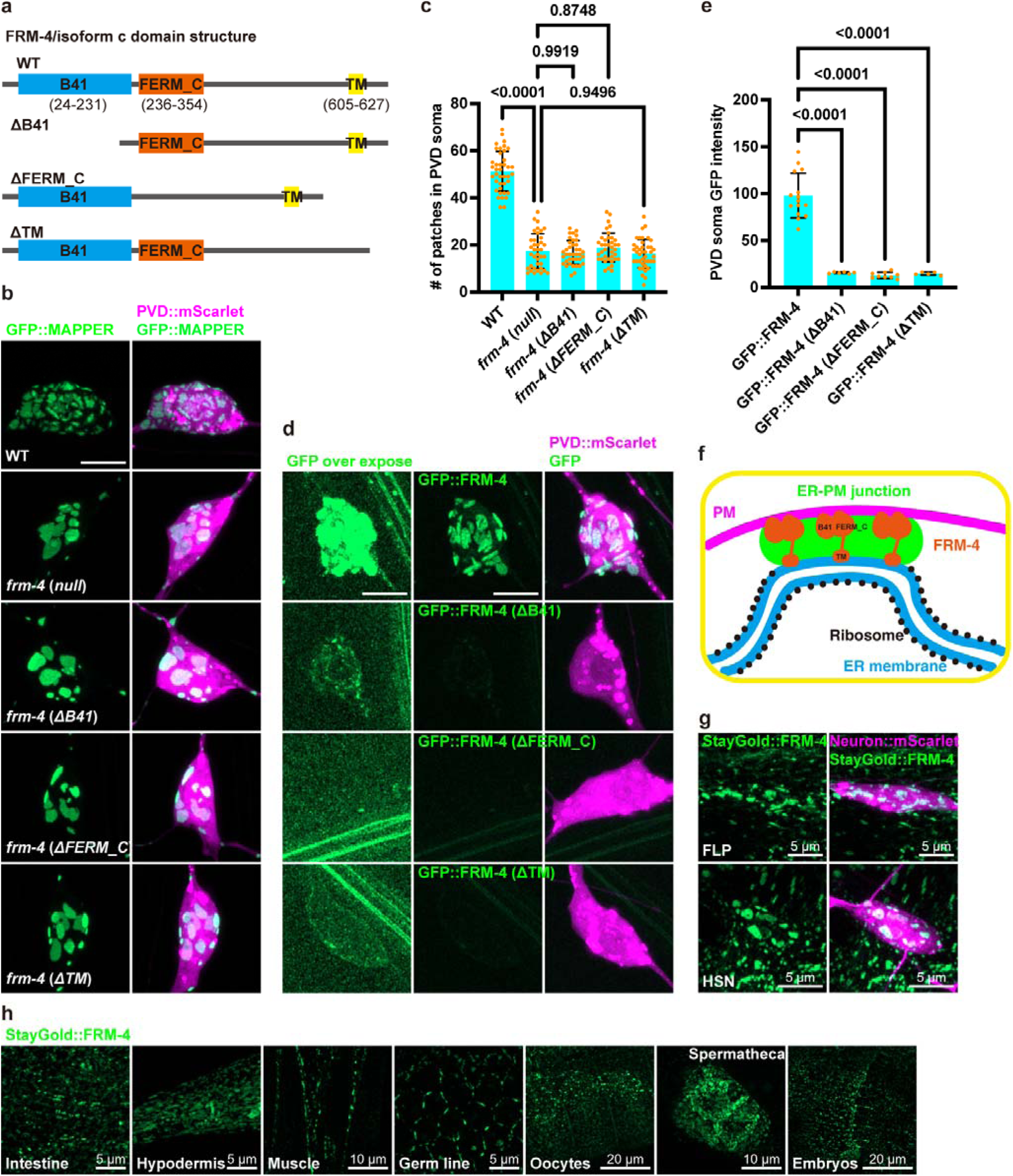
B41, FERM_C and TM domains are all required for FRM-4 regulates EPJ morphology. a, Domain structure of FRM-4, and deletion constructs for the structure-function analyses. b, Representative confocal z-stack images showing EPJs’ pattern in PVD soma in wild type and mutants, visualized by ser-2Prom3::GFP::MAPPER. PVD visualized by ser-2Prom3::mScarlet. Scale bar, 5 µm. c, Quantification the number GFP::MAPPER patches in PVD soma in wild type and mutants. Data are mean ± 95% CI from independent animals. Comparisons using one-way analysis of variance (ANOVA) and Tukey’s tests. d, Representative confocal z-stack images showing endogenous GFP::FRM-4 and domain deletion mutants. PVD visualized by des-2ps::mScarlet. Scale bar, 5 µm. e, Quantification the PVD soma endogenous GFP::FRM-4 and domain deletion mutants GFP intensity. Data are mean ± 95% CI from independent animals. Comparisons using one-way analysis of variance (ANOVA) and Tukey’s tests. f, A schematic model for FRM-4 on EPJ. g, Representative confocal z-stack images of endogenous StayGold::FRM-4 in other neurons soma: FLP and HSN. h, Representative confocal z-stack images of endogenous StayGold::FRM-4 in non-neuronal tissues.

**Extended Data Fig. 3 for Fig. 2.**
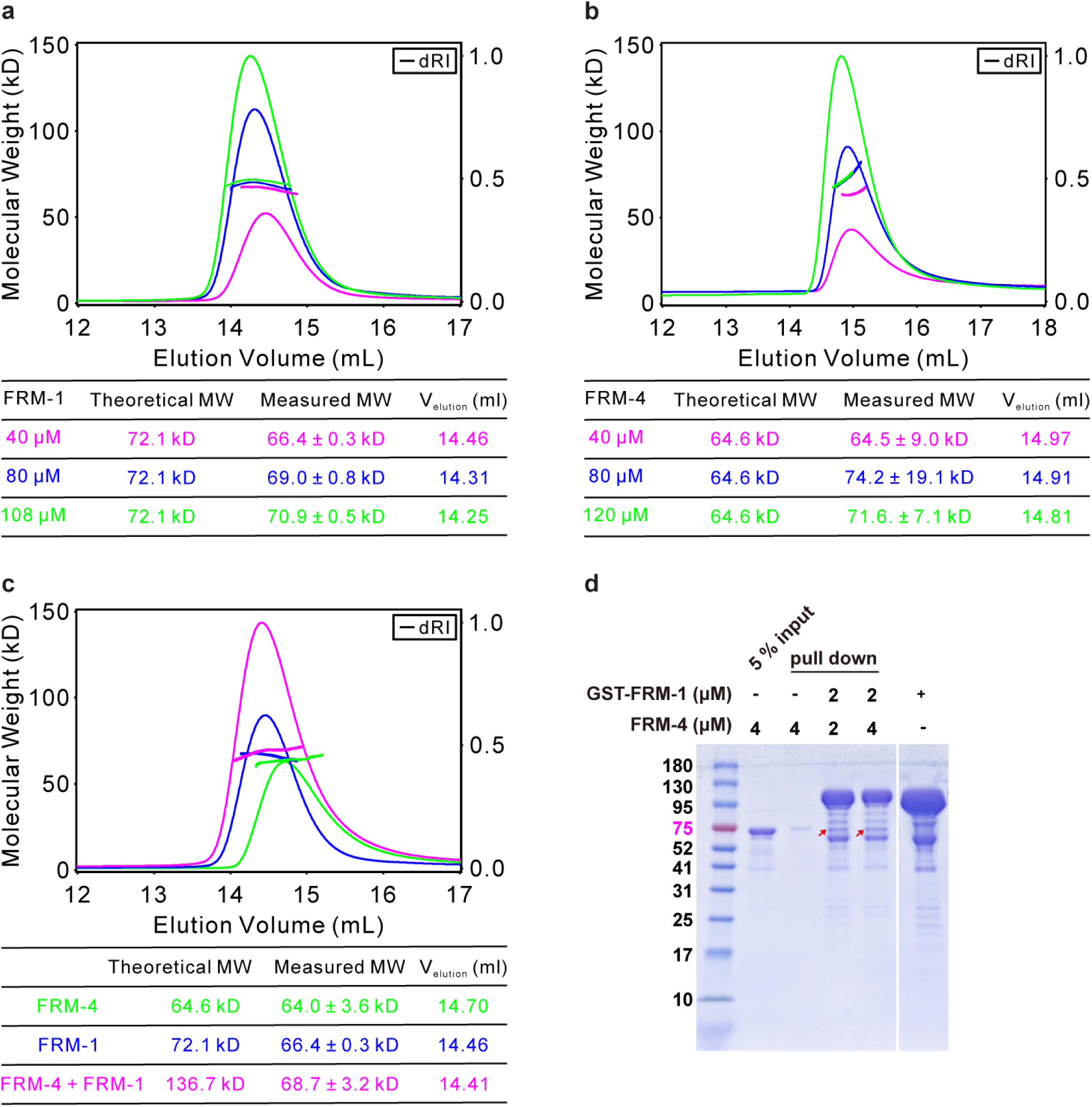
FRM-4, FRM-1 binding in vitro. SEC-MALS analysis of FRM-1 (a), FRM-4 (b). There is a continuous peak shift (Velute) towards smaller elution volume along with increasing protein concentration, indicating concentration-dependent oligomerization of both FRM-1 and FRM-4. c, SEC-MALS detected peak shift towards smaller elution volume upon mixing of FRM-1 and FRM-4, indicating FRM-1 directly binds to FRM-4. d, GST-FRM-1 pull-down assay demonstrated direct binding between FRM-1 and FRM-4. The arrow indicated the FRM-4 band.

**Extended Data Fig. 4 for Fig. 2.**
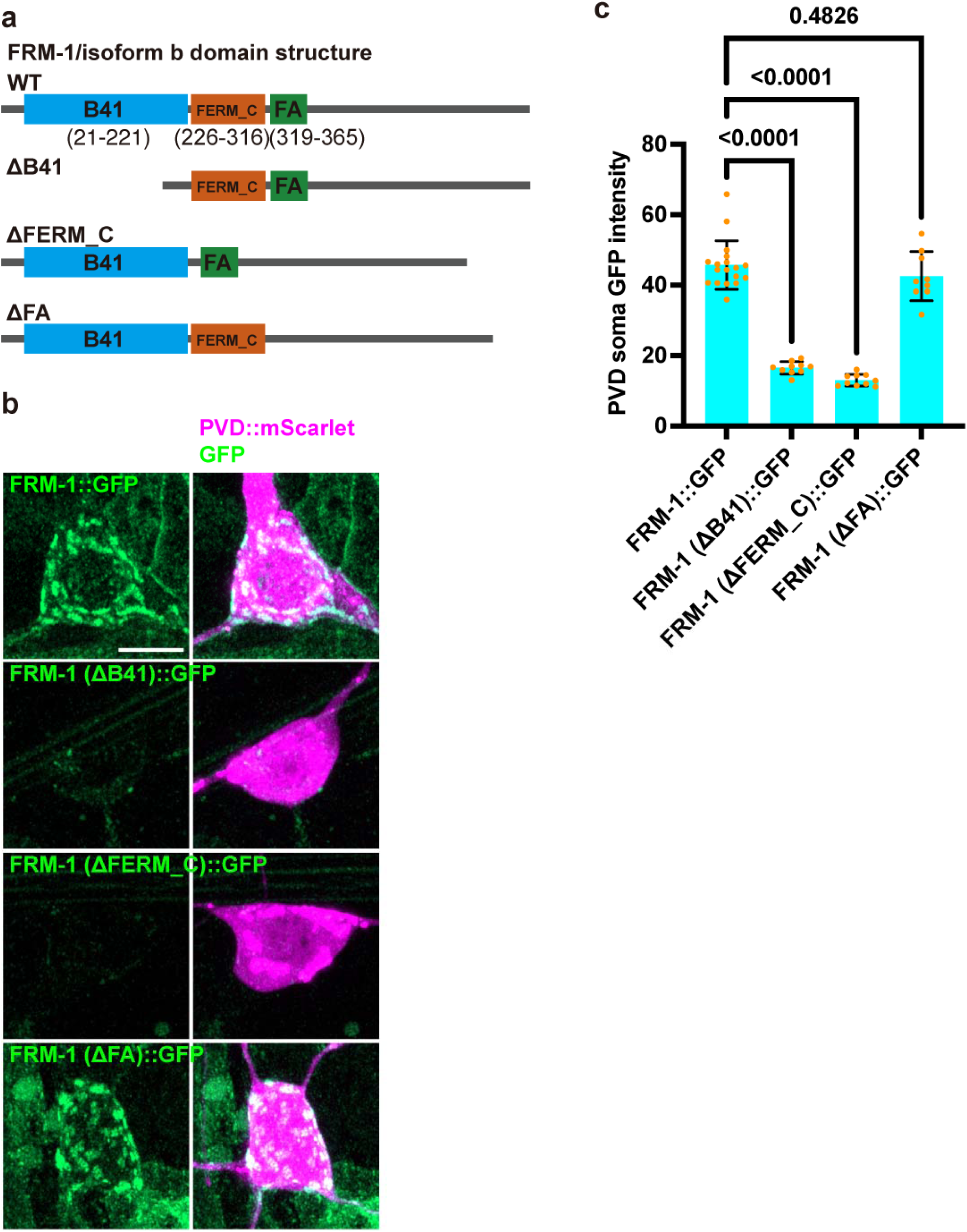
B41, FERM_C domains are all required for FRM-1 protein stabilization. a, domain structure of FRM-1, and deletion constructs. b, Representative confocal z-stack images showing endogenous GFP::FRM-1 and domain deletion mutants. PVD visualized by des-2ps::mScarlet. Scale bar, 5 µm. c, Quantification the PVD soma endogenous GFP::FRM-1 and domain deletion mutants GFP intensity. Data are mean ± 95% CI from independent animals. Comparisons using one-way analysis of variance (ANOVA) and Tukey’s tests.

**Extended Data Fig. 5 for Fig. 4.**
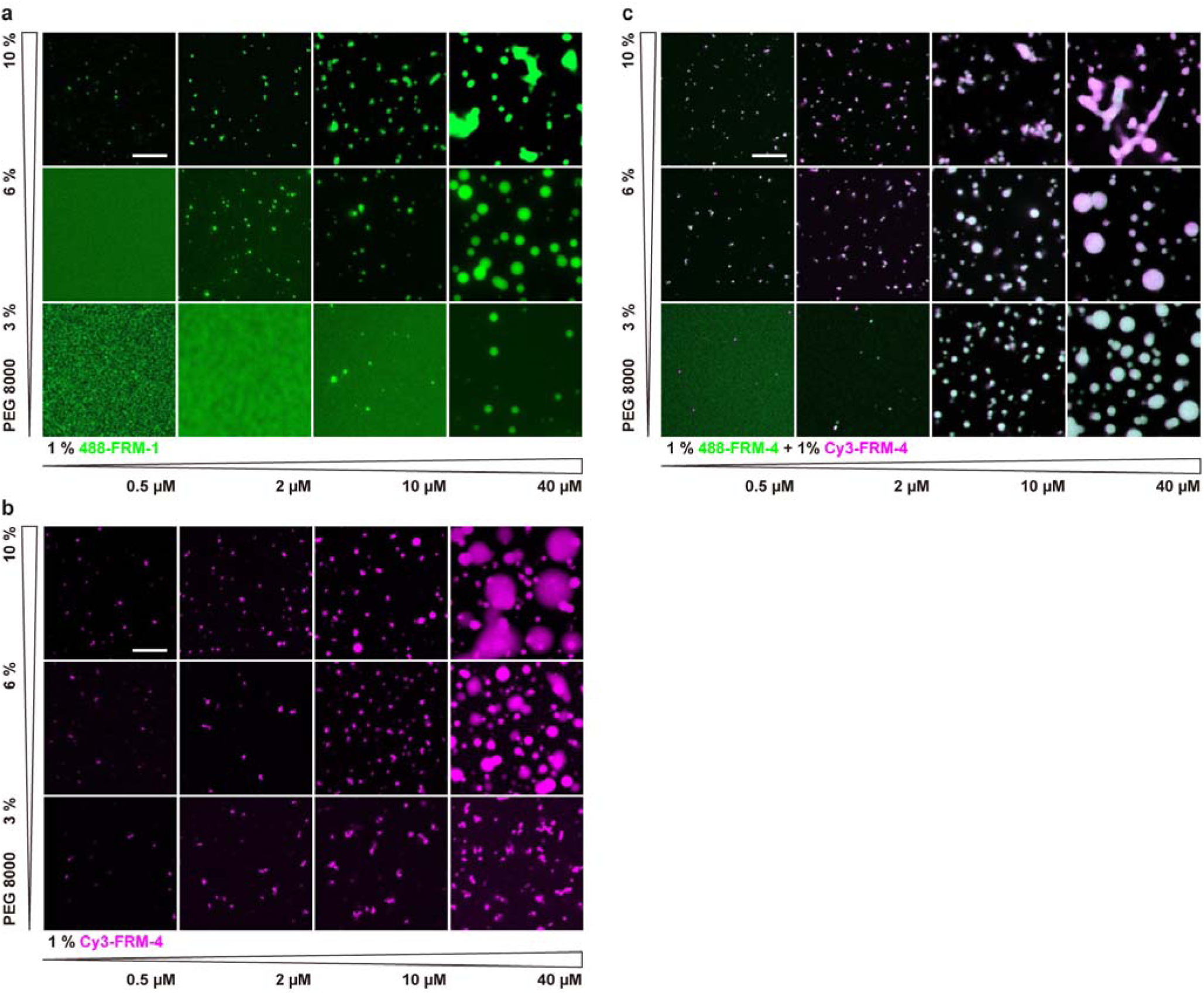
FRM4, FRM1 form condensates. Phase diagram of FRM1 (a), FRM4 (b), and FRM1/FRM4 complex (c). The higher the protein or PEG concentration, the stronger the phase separation. Too high concentration of PEG may modify the condensate rigidity. Scale bar, 10 µm.

